# Loss of intermediate filament IFB-1 reduces mobility, density and physiological function of mitochondria in *C. elegans* sensory neurons

**DOI:** 10.1101/723205

**Authors:** Syed Nooruzuha Barmaver, Muniesh Muthaiyan Shanmugam, Yen Chang, Prerana Bhan, Gong-Her Wu, Oliver I. Wagner

**Author notes:** Both authors contributed equally to this study (Syed Nooruzuha Barmaver, Muniesh Muthaiyan Shanmugam). **Corresponding author**: Dr. Oliver I. Wagner, Professor, National Tsing Hua University, Institute of Molecular and Cellular Biology & Department of Life Science, 101, Sec. 2, Kuang Fu Road, Hsinchu 30013, Taiwan (R. O. C.).

## Abstract

Mitochondria and intermediate filament (IF) accumulations often occur during imbalanced axonal transport leading to various types of neurological diseases. It is still poorly understood whether a link between neuronal IFs and mitochondrial mobility exist. In *C. elegans*, among the 11 cytoplasmic IF family proteins, IFB-1 is of particular interest as it is expressed in a subset of sensory neurons. Depletion of IFB-1 leads to mild dye-filling and significant chemotaxis defects as well as reduced life span. Sensory neuron development is affected and mitochondria transport is slowed down leading to reduced densities of these organelles. Mitochondria tend to cluster in neurons of IFB-1 mutants likely dependent on fission but independent of fusion events. Oxygen consumption and mitochondrial membrane potential is measurably reduced in worms carrying mutations in the ifb-1 gene. Membrane potential also seems to play a role in transport such as FCCP treatment led to increased directional switching of mitochondria. Mitochondria colocalize with IFB-1 in worm neurons and appear in a complex with IFB-1 in pull-down assays. In summary, we propose a model in which neuronal intermediate filaments may serve as critical (transient) anchor points for mitochondria during their long-range transport in neurons for steady and balanced transport.

**Abstract Figure:** 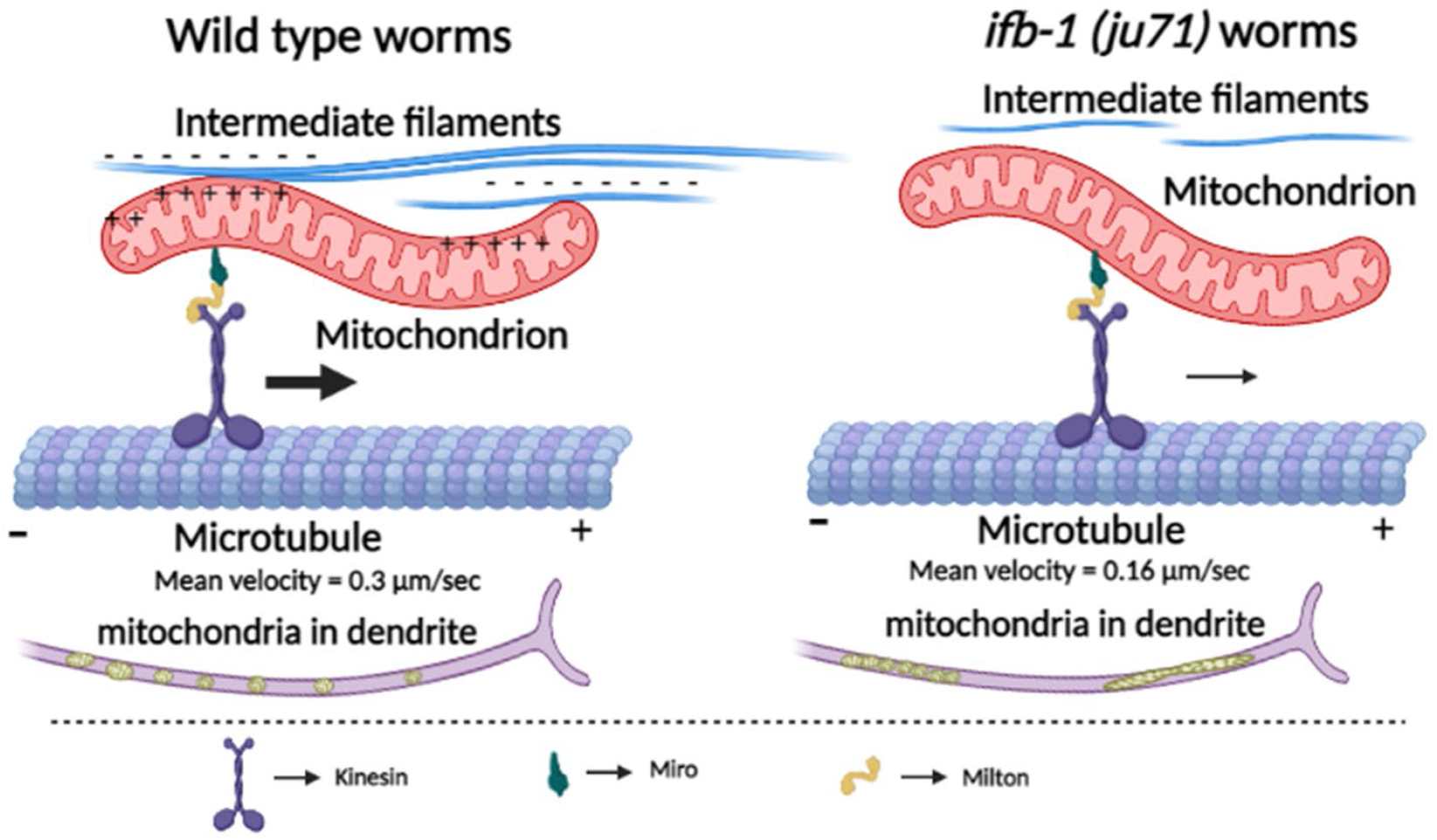

**Synopsis:** Various neurological diseases are both associated with abnormal accumulations of neuronal intermediate filaments as well as mitochondria. Here, we report a link between these two phenomena employing the model organisms *C. elegans*. Depletion of neuronal intermediate filament IFB-1 impairs the transport of mitochondria in sensory neurons leading to clustered and accumulated mitochondria affecting neuronal growth and oxygen consumption in nematodes.

## INTRODUCTION

The neuronal cytoskeleton is composed of three major cytoskeletal elements: actin, microtubules and intermediate filaments (IFs). IFs are divided into six classes based on their sequence homologies, and in mammalian neurons, a diverse set of IFs can be found, including synemin, peripherin, vimentin, nestin, α-internexin and neurofilaments (NFs). Abnormal accumulations of NFs is found in various neurological diseases such as Parkinson’s disease (PD), amyotrophic lateral sclerosis (ALS), infantile spinal muscular atrophy (SMA), Alzheimer’s disease (AD), Charcot-Marie-tooth (CMT) disease, Giant axonal neuropathy (GAN) and hereditary sensory-motor neuropathy (HSMN); while many of the aforementioned IFs (specifically peripherin and α-internexin) are known to be central components of NFs ^1–3^. IFs act as a molecular scaffold and regulate intracellular positioning of nucleus and organelles providing integrity to cells and tissues ^4^. A study demonstrates that IFs alter the dynamics of microtubules to affect remodeling of developing synapses and that IF accumulations hyperstabilize microtubules ^5^. Mitochondrial quality control, membrane potential, organization and function are all impaired upon loss of IFs ^6–8^. It has also been shown that mutations in IFs affect mitochondrial size, density and respiratory function ^8–11^.

In *C. elegans*, the IF family is composed of eleven IF proteins that can be divided into two categories based on their expression patterns: IFB-2, IFC-1, IFC-2, IFD-1, IFD-2, and IFP-1 belong to one category of IF proteins that localize to the intestinal terminal web where they provide mechanical flexibility to the intestinal lumen. The other category includes IFA-1, IFA-2 (previously named MUA-6), IFA-3, IFA-4 and IFB-1. In this category, IFB-1 co-expresses and forms obligate heteropolymers with either one of the IFAs. These IFs are widely expressed in worms and except IFA-4, all are essential for viability ^12–14^. Among these IFs, IFA-1, IFA-2, and IFB-1 particularly attracted our attention as they are expressed in *C. elegans* amphid neurons ^15^. Since IFB-1 forms heteropolymers with IFA-1 and IFA-2 ^13^, we focused on the essential IFB-1 and its presumed effects on mitochondrial transport in amphid sensory neuronal bundles. To monitor mitochondria in the amphid sensory neuron bundle and to overexpress IFB-1 for colocalization experiments, we employed the *Posm-5* promoter to drive the expression of genes in amphid, labial and phasmid sensory neurons ^16,17^.

As a result of alternative splicing, IFB-1 can be found in two isoforms, IFB-1A and IFB-1B. IFB-1A and IFB-1B transcripts comprise different promoters and distinct first exons, but share the remaining C-terminal domains (Fig. 1N); thus, IFB-1A and IFB-1B differ in their N-terminal head domains. Although two IFB-1 isoforms are slightly different in their expression sites and functions, they are both required for muscle attachment and epidermal elongation at the 2-fold stage in *C. elegans* embryonic development. Both IFB-1 isoforms localize to epidermal attachment structures linking muscles to cuticles. An antibody raised against the C-terminal region of IFB-1 does detect both isoforms of IFB-1, whereas in worms carrying the *ju71* mutant allele (which removes the promoter region of the *ifb-1* gene) only the isoform IFB-1B can be detected by Western blots ^18^. Another allele *ok1317* exist, but has not been characterized yet. Our analysis reveals that it carries a 1361 bp deletion in the promotor region, 1032 bp upstream of the start codon of *ifb-1a* (and both *ju71* and *ok1317* located in the first intron of *ifb-1b*) (Fig. 1N).

**Figure 1:**
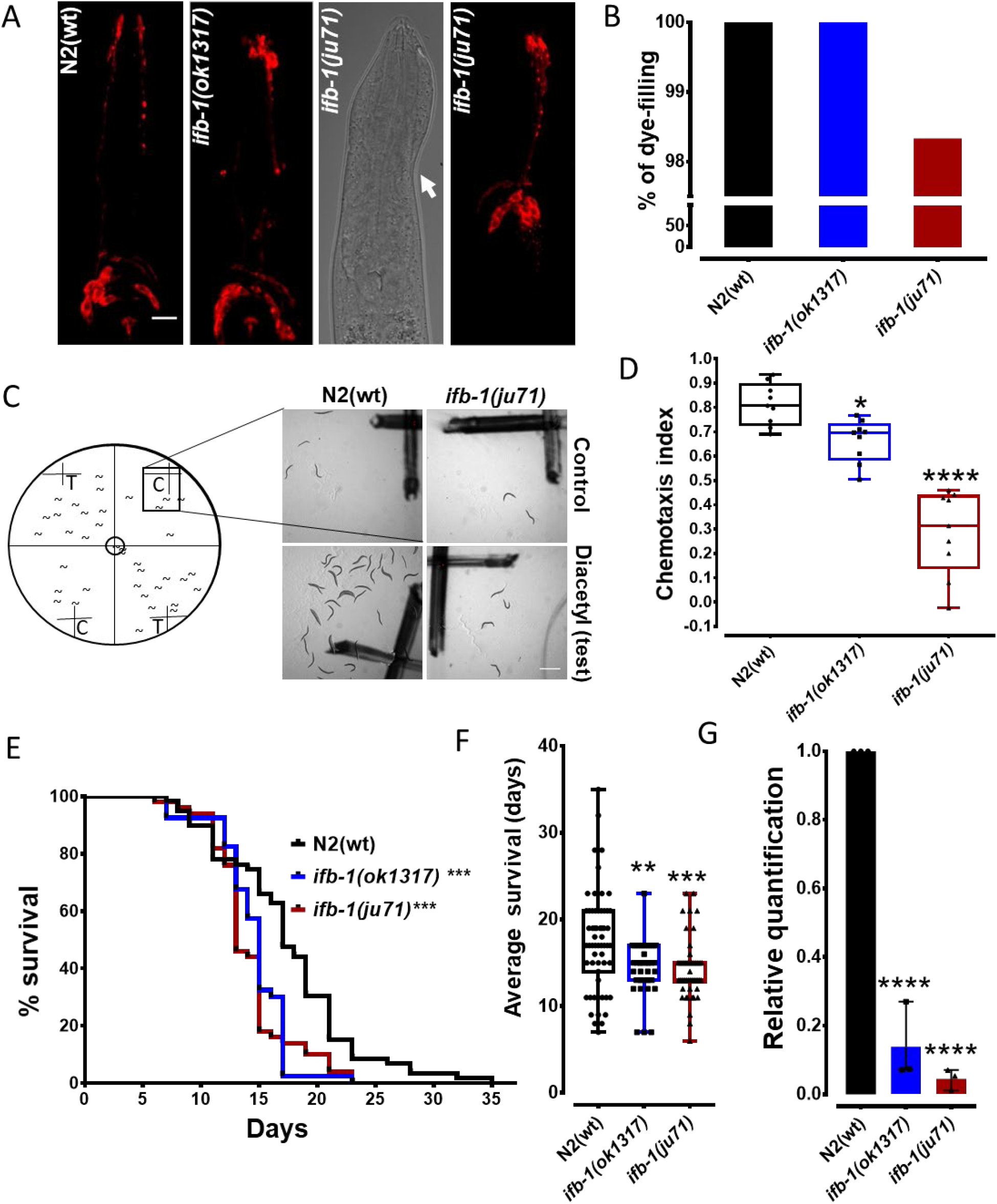
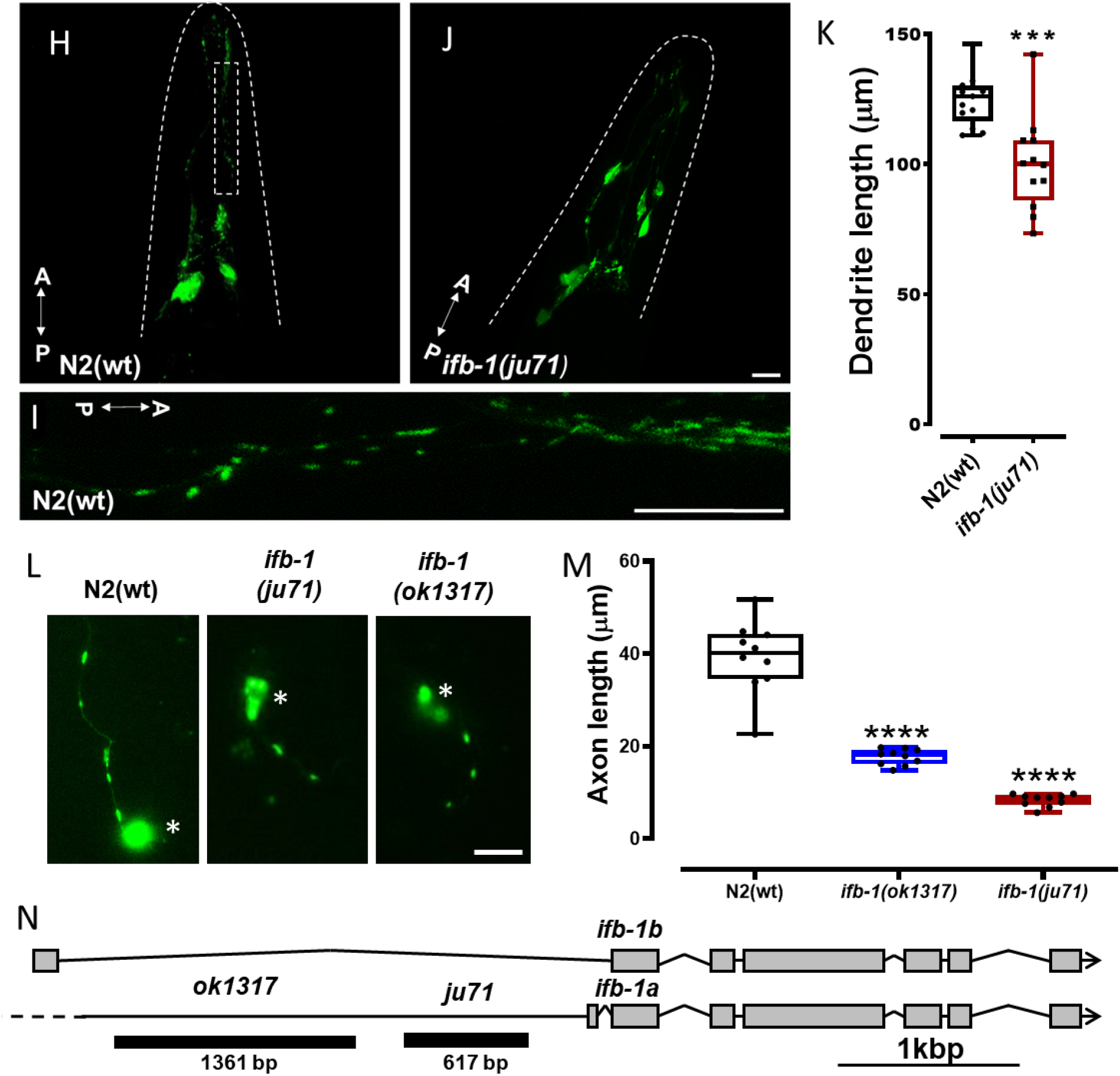
Characterization of *ifb-1* mutants. (A) Representative images of ciliated endings of amphid dendrites at the nose of various worms (N2 control, *ok1317* and *ju71* mutants) after dye-filling. Arrow indicates structural deformities in *ju71* mutant (hook-head phenotype). (B) Dye-filling quantification (N = 300 worms). (C) Pictorial representation (left panel) and photograph of representative plates (right panel) of quadrant chemotaxis assay. (D) Chemotaxis index from different genetic backgrounds (N = 30-150 worms with 9 trials). (E) Percentage survival rate (N = 40-59 worms) and (F) average survival days of worms from different genetic backgrounds. (G) Quantitative PCR analysis of *ifb-1* mRNA expression. (H) Confocal images of N2 worms expressing *Posm-5::mito::gfp*. Dashed outlines depict the margins of worm’s heads. (I) Mitochondria in amphids magnified from the dashed rectangle in (H). (J) *ifb-1*(*ju71)* mutant expressing *Posm-5::mito::gfp*. (K) Quantification of dendrite lengths measured from the somas of amphid neurons (located in the nerve ring) to distal regions of the dendrite (where cilia begin to emanate to the nose) (N = 12-13 worms). (L) Representative images of cultured worm neurons from different genetic backgrounds. (M) Quantification of axonal length from neurons shown in (L) (N = 30 cells). (N) Graphical representation of ifb-1 exon/intron gene structure and the revealed location of the deletion mutations. Scale bars: 10 μm in (A, H-J and L) and 1 mm in (C). Error bars: ± max. and min. range. One-way ANOVA with Fisher’s LSD test (D, F, G), Student’s t-test (K) and Gehan-Breslow-Wilcoxon test (E). *p<0.05, **p<0.01, ***p<0.001 and ****p<0.0001. A – anterior direction and P – posterior direction.

Mitochondria produce ATP via the oxidative phosphorylation chain, whereas a toxic by product, ROS (reactive oxygen species), is known to damage cellular components and mtDNA. Mutations in mtDNA are accumulated over lifetime resulting in reduced mitochondrial function with aging. In many neurodegenerative diseases (with impaired axonal transport) mitochondria appear largely damaged. For example, mutations in the mitochondrial proteins PINK1, mitofusin 2 and paraplegin are causatives of Parkinson’s disease. In HSP, CMT and in neurons with mutant forms of SOD1, parkin, huntingtin, PARK1/DJ-1 all damage mitochondria. Amyloid-β-peptide, the neurotoxin MPP+ and excitotoxic glutamate damage mitochondria, and at the same time impair axonal transport; resulting in increased net retrograde movements of mitochondria with the consequence that these organelles accumulate in somas. Axonal depletion of mitochondria and accumulations in somas can be seen in human ALS cases, therefore mitochondria are not only damaged in mutant SOD1 axons but also their number in axons is severely reduced ^19–21^.

The current understanding of mitochondrial transport is that conventional kinesin-1 (KIF5) and dynein are attached at the same time to a mitochondrion, occasionally pulling in opposite directions, thus causing bidirectional and saltatory movements of these organelles. Recently, adaptor proteins have been described to facilitate binding of KIF5 to mitochondria (syntabulin or miro/milton) revealing previous unknown complexity of motor-organelle recognition ^22–24^. Interactions with the neuronal actin cytoskeleton (axonal cortex and at growth cones) occasionally keep mitochondria stationary (only about one third of axonal mitochondria are mobile) ^25^. Similarly, interactions between mitochondria and NFs have been reported *in vitro* ^26^; however whether such interactions occur *in vivo* is not known. Notably, SNPH (syntaphilin) directly docks (cross-links) mitochondria to microtubules ^27^. The importance of immobilizing mitochondria lies in their localized distribution depending on cellular ATP needs. Indeed, mitochondria accumulate at those areas where ATP (and calcium) consumption is high ^28,29^. Based on the thin diameter of the axon and its crowded internal ^30^, it is apparent that large organelles such as mitochondria may reside in close proximity to cytoskeletal elements other than microtubules. These filamentous elements include actin and IFs that might act as static anchor-points to immobilize mitochondria ^31^. Critically, in non-neuronal cells, IF proteins are reported to affect mitochondrial morphology, distribution and motility. For example, it has been shown that vimentin directly binds to mitochondria to affect mitochondrial morphology, organization and mobility in fibroblasts ^32,33^. Further, studies have shown that mitochondrial motility is significantly diminished upon accumulations of IFs ^10,11^ and in neurons, the transport of NFs depends on kinesin-1 ^34^. It has been also revealed that IF-associated protein plectin 1b links mitochondria to the IF-network in fibroblasts ^35^ and that desmin, a type III IF, directly interacts with mitochondria through its N-terminus ^9^.

Because the role of neuronal IFs on axonal transport has been poorly investigated, we set out to understand whether mutations in IFB-1 (one of the few IFs expressing in *C. elegans* neurons) would affect mitochondria transport. The significance of this research is that both neuronal IF and mitochondria accumulations are a hallmark of a large number of human neurological diseases.

## RESULTS

Above, we cited several reports demonstrating direct IF-mitochondria interactions as well as the effect of IFs on the mobility of mitochondria; however, most of these studies employ cell culture systems. Also, though NF-mitochondria interactions have been dissected *in vitro* ^26^, their *in vivo* interactions remain to be demonstrated. Though recently we identified and characterized a NF ortholog NEFR-1 (TAG-63) in *C. elegans* ^36^, several IFs are expressed in amphids and phasmids sensory neurons of *C. elegans* ^12^. Specifically, IFB-1 has been shown to express in amphid neuron bundles ^15^. Importantly, the allele *ju71* carries a deletion in the promoter region of the *ifb-1* gene (Fig. 1N) resulting in a reduced expression of IFB-1A (but not IFB-1B) in *C. elegans* as revealed by Western blots ^18,37^ and quantitative PCR (Fig. 1G). Because *ifb-1* is embryonic lethal ^13,15^ expression of IFB-1B ^18,37^ in *ju71* is beneficial for the animal’s survival (though strong phenotypes such as uneven epidermis and “hook head” are obvious, Fig. 1A). In addition to the *ju71* allele, another allele *ok1317* exists that carries a 1361 bp deletion in the promoter region (upstream of the *ju71* deletion) (Fig. 1N). Though dye filling (as a measure for cilia structure impedances) is not affected in *ok1317* mutants, mild chemotaxis defects can be detected (Fig. 1B and D). Other phenotypic parameters such as life span and axonal length development (primary cultures from *C. elegans* embryos) are strongly reduced in *ok1317* (similarly as observed for the *ju71* background) (Fig. 1E+F and M). We then designed a plasmid that drives the expression of *mito::gfp* (a C-terminal GFP fused to an N-terminal mitochondria-targeting sequence) in amphid neurons employing the promoter *Posm-5* ^16^. After microinjecting the plasmid, resulting strain *N2;nthEx4[Posm-5::mito::gfp;rol-6(su1006)]* (Fig. 1H+I) was then crossed with either CZ4380 *ifb-1(ju71)* or RB1253 *ifb-1(ok1317)* strains, and genotyped to confirm homozygosity. Positioning worms with their vulva sidewise, amphid neurons can be clearly distinguished from labials as amphids seem to adopt the shape of the pharynx; and labials are located closer to the (lateral) cuticle of the worm. The somehow shortened and thickened head in the *ju71* background obviously affects morphologies of amphid neuron bundles which noticeably became retracted (Fig. 1A) that may explain defects in chemotaxis (Fig. 1D) ^38^.

We then used Nipkow spinning disk microscopy to track and analyze the motility of mitochondria in amphid neuron bundles. Single mitochondria can be clearly observed in the amphid bundle (Fig. 1I); however, based on resolution limitations, mitochondria in single neurites of these bundles (encompassing 12 tightly packed neurites) cannot be discriminated. For motility assays, we determined anterograde (Fig. 2A) and retrograde (Fig. 2B) velocities (without pauses), the occurrences of directional changes (mitochondria reversals, Fig. 2C) as well as number of pauses in a defined region of the neuron (Fig. 2D). In general, effects are more pronounced in *ju71* mutants as opposed to the *ok1317* allele (Fig. 2) which is in accordance with the milder observed phenotypes in *ok1317* backgrounds (Fig. 1). Based on these results, we also performed rescue experiments only in *ju71* mutants (Fig. 2). Though pausing of mitochondria is not affected in *ju71* worms, velocities and mitochondrial reversals are both significantly affected in *ju71* (Fig. 2A-D). Because mitochondrial motility is significantly affected in *ifb-1* mutants, we asked whether this effect would result in abnormal mitochondrial distribution patterns in amphids. Here, we analyzed changes in densities along the neurites, Feret’s diameter (“maximum diameter”) and size (area) (Fig. 3A, C-E). As a result, mitochondria appear larger (likely clustered) in the neurites of *ju71* mutants with concomitantly reduced densities (Fig. 3 A, C-E). Though in *ok1317* backgrounds mitochondria densities are decreased, their sizes remain unaffected (Fig. 3 A, C-E). To investigate whether the determined size changes of mitochondria in *ju71* mutants are caused by the fusion and fission machinery, we knocked down fission gene fis-1 and fusion gene eat-3 in wild type and *ju71* mutants (Fig. 3B-E). Knocking down fis-1 in N2 worms leads to increased mitochondrial sizes as expected, such as fission events are suppressed (Fig. 3C). Similarly, knocking down eat-3 in N2 leads to significantly decreased sizes as expected, since fusion events are suppressed. Both fis-1 and eat-3 did not further affect the *ju71* phenotype (Fig. 3C). An increase in Feret’s diameter is usually the result of more elongated mitochondria in a population. Indeed, knocking down fis-1 gene in N2 leads to markedly increased Feret’s diameters (Fig. 3D). Even in *ju71* mutants fis-1 knockdown leads to an additional increase in Feret’s diameter. As expected, knocking down eat-3 in N2 worms results in smaller Feret’s diameters, and this knockdown did not further affect the *ju71* phenotype (Fig. 3D). Mitochondrial density is supposed to decrease when knocking down fis-1 due to expectedly overall larger mitochondria in a defined area of the neurite. At the same time knocking down eat-3 should lead to more fragmented mitochondria increasing densities in the neurites. Indeed, knocking down fis-1 in N2 results in decreased mitochondrial densities; however, eat-3 knockdown lead to comparable densities as seen in wild types (Fig. 3E). Nevertheless, also here (similar to effects seen on Feret’s diameter), fis-1 knockdown did further affect the *ju71* phenotype while eat-3 did not change *ju71* the phenotype (Fig. 3E).

**Figure 2:**
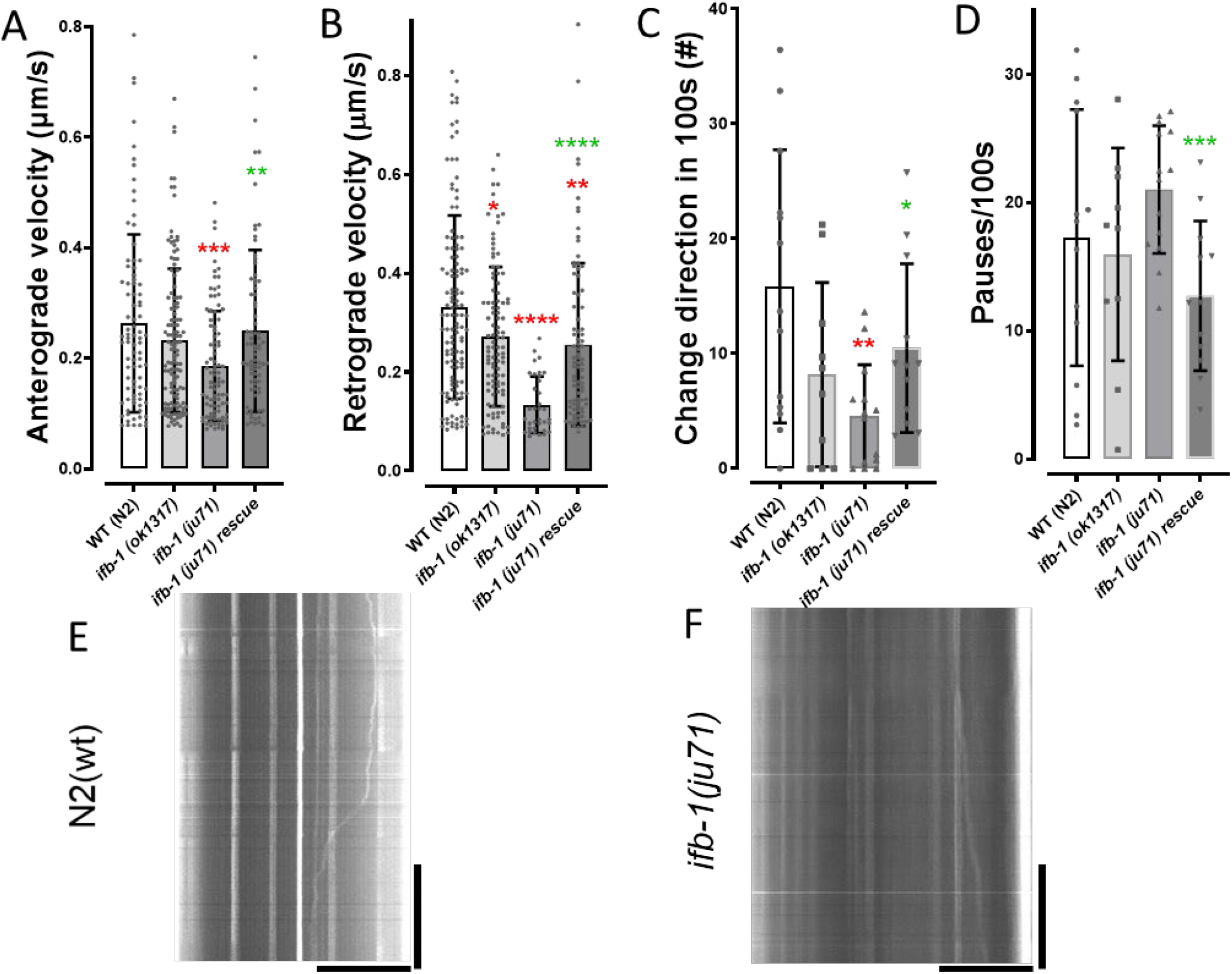
Motility analysis of mitochondria in amphid neuron bundles of living *C. elegans*. (A) Anterograde and (B) retrograde velocities in wt (referred to strain *N2;nthEx4[Posm-5::mito::gfp;rol-6(su1006)]*) and different mutant backgrounds including rescue experiments. (C) Directional changes (mitochondria reversals) in wt and mutants. (D) Quantification of pausing behavior in wt and mutants. (E-F) Representative kymographs of mitochondria movements in wt and *ju71* mutants. Total number of analyzed moving events: 381 for wt, 382 for *ifb-1(ju71)*, 278 for *ifb-1(ok1317)*, and 285 for *ifb-1(ju71)* rescue. One-way ANOVA with Dunnett’s test for multiple comparisons. **p<0.005, ***p< 0.001, ****p<0.0001. Green asterisk denotes significant differences between *ifb-1*(*ju71*) and *ifb-1*(*ju71*) rescue strain using unpaired Student’s t-test. Error bars: ± SD. Horizontal scale bar: 20 μm. Vertical scale bar: 40 s.

**Figure 3:**
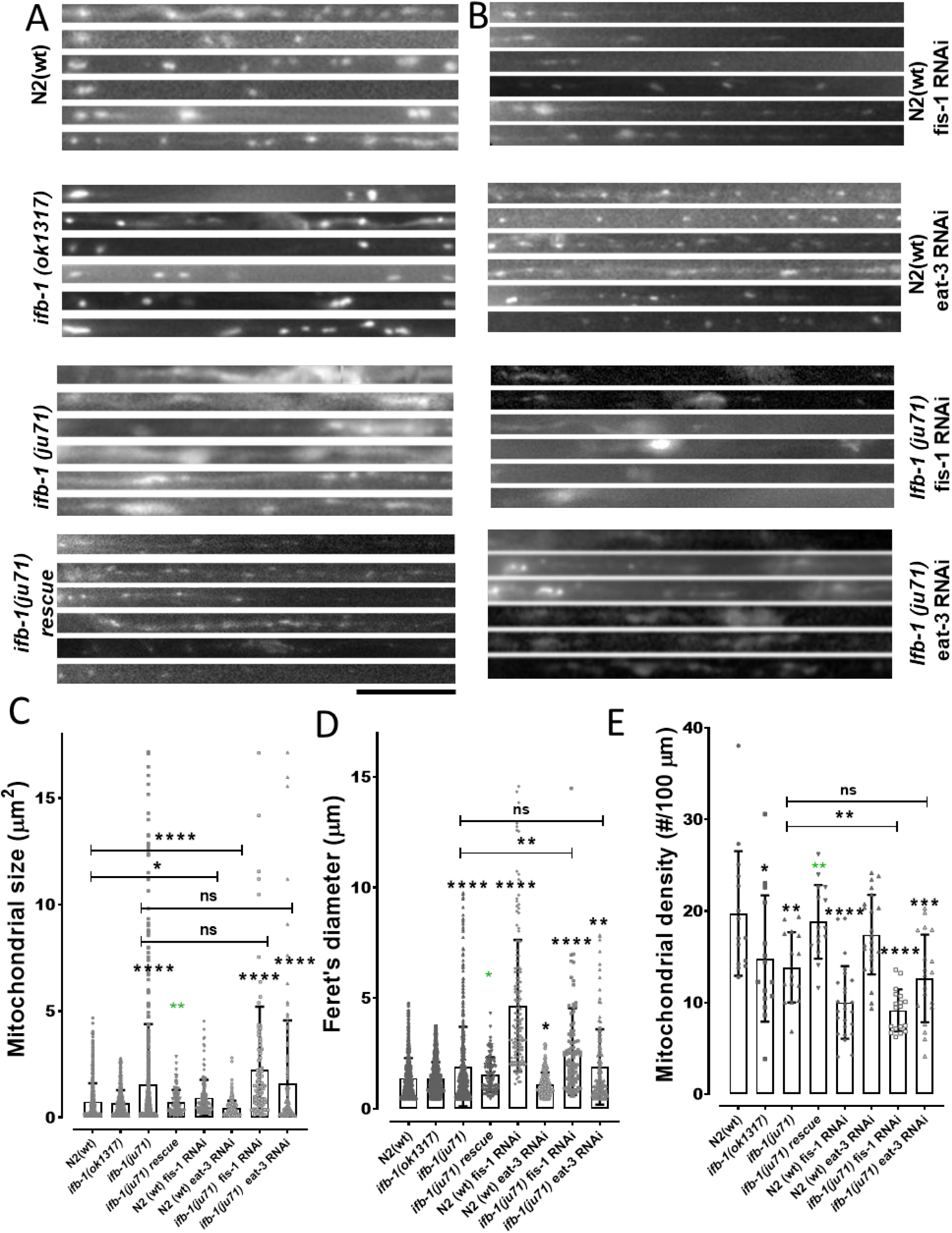
Mitochondrial morphologies and densities in amphid neurons in *ifb-1* mutants and dependencies on genes fis-1 and eat-3. (A-B) Straightened and stacked amphid dendrites from different genetic backgrounds. (C) Quantification of mitochondrial sizes in these dendrites. (D) Quantification of Feret’s diameter from images in (A-B). (E) Density quantification of mitochondria. Scale bar: 10 μm. Error bars: ± SD max. and min. range. N = 559-841 particles in (D+E) and n = 15-25 dendrites in (C). One-way ANOVA with Dunnett’s test for multiple comparisons. **p<0.005, ***p< 0.001, ****p<0.0001. Green asterisk denotes significant differences between *ifb-1*(*ju71*) and *ifb-1*(*ju71*) rescue strain using unpaired Student’s t-test.

Reduced mobility of mitochondria (Fig. 2) and increased clustering (Fig. 3) has likely physiological consequences for the neuron. Indeed, neurites of amphids are shorter in *ju71* worms (Fig. 1K) as well as axons appear shorter in cultured primary neurons from *ju71* worms (Fig. 1M). To understand whether reduced mobility and clustering of mitochondria could be based on secondary effects resulting from ifb-1 gene depletion (such as affecting the physiological function of mitochondria), we wanted to know if mitochondria mobility is generally affected in mitochondria at low physiological states. To induce such states, we employed the uncoupling agent FCCP (carbonyl cyanide p-(trifluoromethoxy)phenylhydrazone) which is an ionophore that reduces mitochondrial membrane potential and concomitantly ATP production. Because uptake efficiency of FCCP in living nematodes is not well studied, we used cultured primary neurons (from *C. elegans* embryos) that are more likely to efficiently uptake FCCP. Interestingly, FCCP (1 μM) affected mitochondria reversals (Fig. 4C) as well as increased the percentage of stationary mitochondria (Fig. 4F), but did not affect their transportation speeds (Fig. 4A+B). We also attempted to observe mitochondrial motility changes in neuronal cultures from *ju71* worms, however, effects are only mild and most likely due to the fact that these cultures are a mixture of all neurons, and amphids (as well as phasmids) only make out a fraction of these.

**Figure 4:**
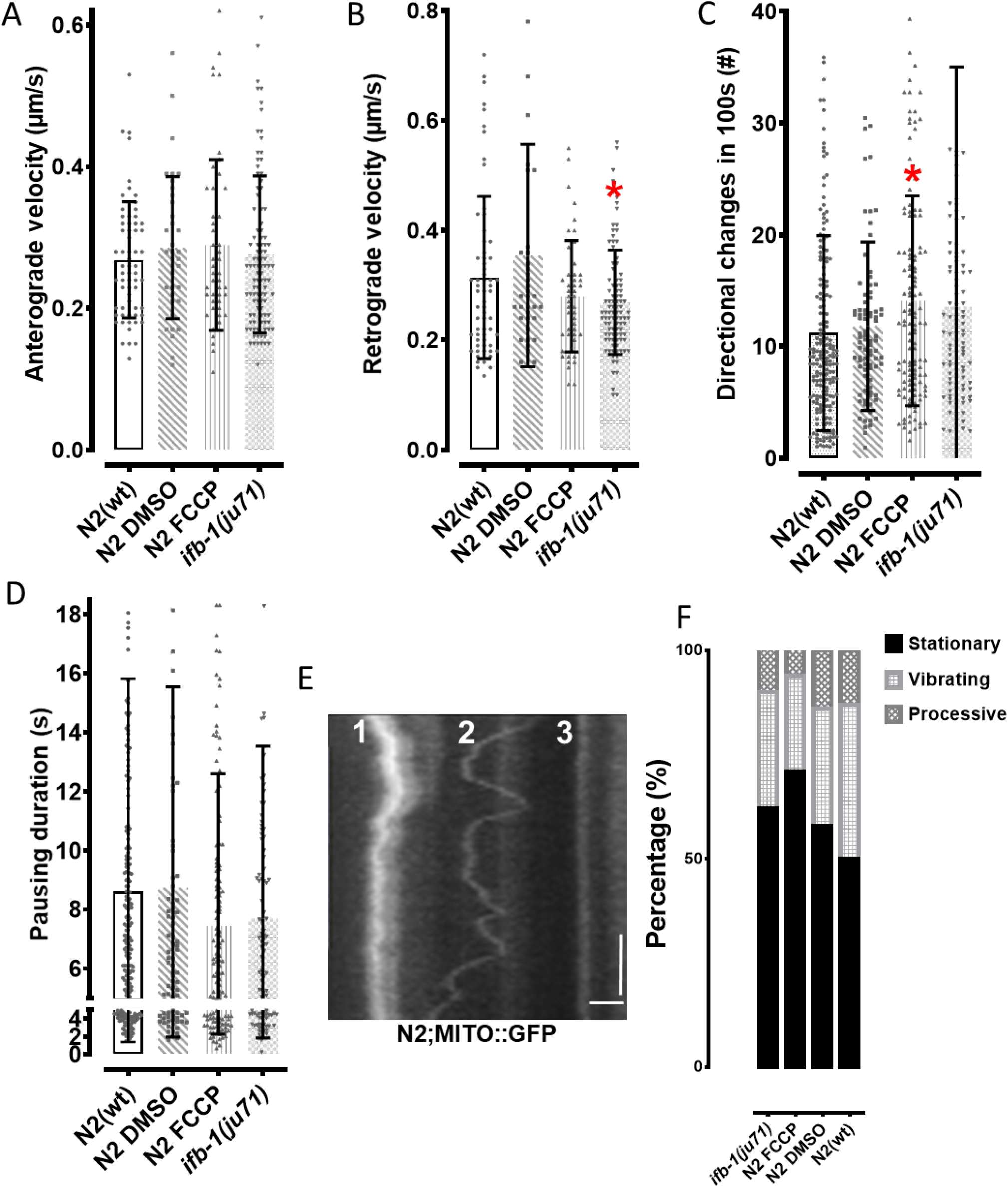
Motility analysis of mitochondria in neuronal cultures before and after 1 μM FCCP treatment. Quantification of (A) anterograde and (B) retrograde velocities, (C) directional changes, and (D) pausing durations. (E) Example of kymograph depicting vibrating (1), processive (2) and stationary (3) mitochondria (left panel), and (F) percentage of mitochondria from these different motility states (right panel) (F). Total number of analyzed moving events: 3697 for wt and 2217 for *ifb-1(ju71)*. Horizontal scale bar: 10 μm. Vertical scale bar: 20 s. Error bars: ± SD. One-way ANOVA with Dunnett’s test for multiple comparisons. *p<0.05,. **p<0.005, ***p< 0.001, ****p<0.0001.

Though the physiological state may only partially affect the transportation of mitochondria (such as increased reversals, Fig. 4C), we wanted to know more about the physiological consequences resulting from mitochondrial accumulations and reduced transport efficiency upon ifb-1 depletion in worms. Here, we employed the XF Extracellular Flux Analyzer (Seahorse Bioscience) that allows for the determination of oxygen consumption of living *C. elegans* animals in solution. Interestingly, oxygen consumption of nematodes is generally reduced in *ju71* backgrounds (Fig. 5C+D). Moreover, using the mitochondrial membrane potential sensitive dye TMRE, we detected reduced mitochondrial membrane potential in *ju71* mutants (Fig. 5E+F). However, from Figure 4 we can discern that membrane potential may not be the deciding factor for reduced transportation based on ifb-1 depletion.

**Figure 5:**
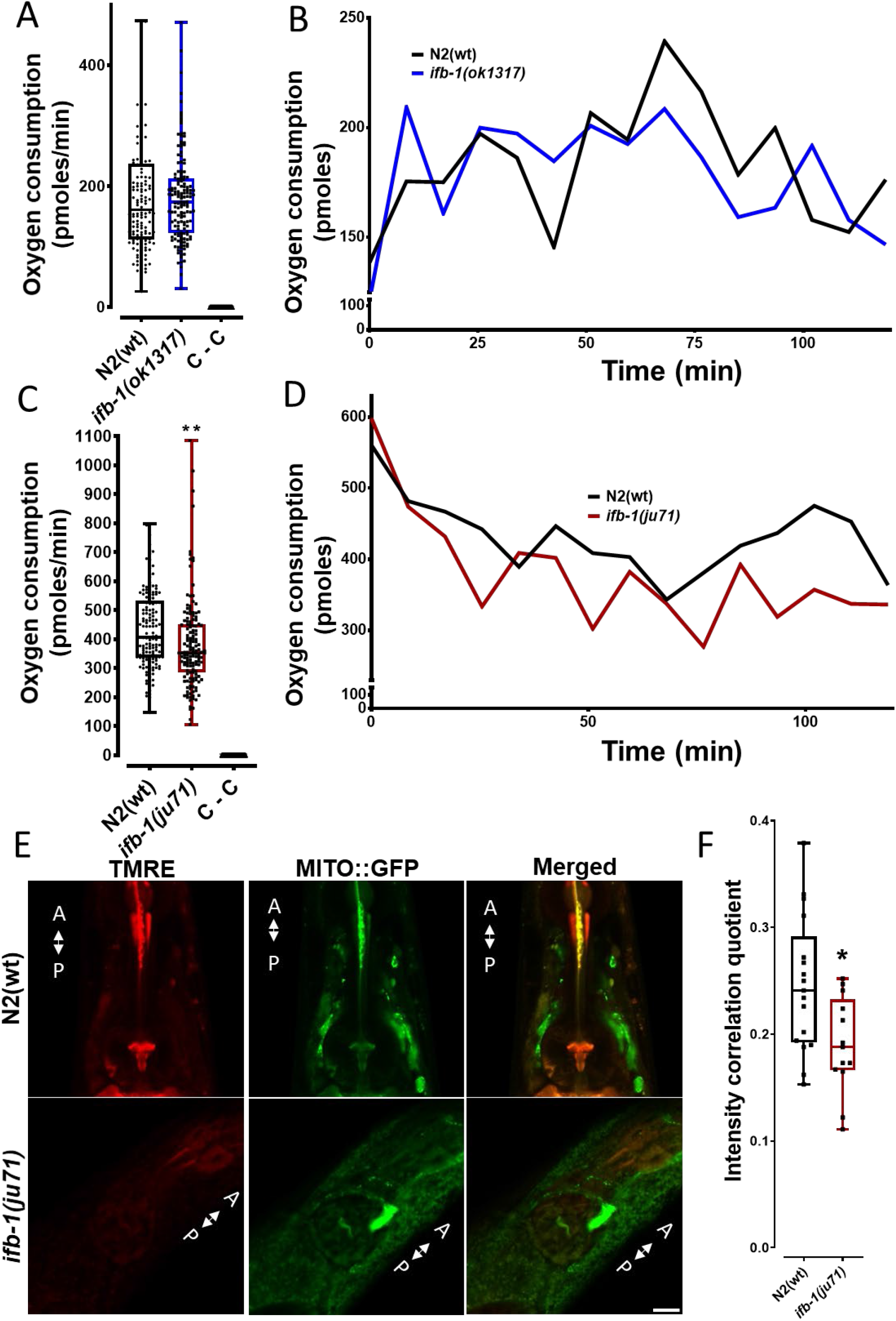
Oxygen consumption of wild type and ifb-1 mutant worms. (A) Average oxygen consumption of N2 control worms and *ifb-1(ok1317)* mutants. (B) Oxygen consumption over the assayed period of 120 min. (C) Oxygen consumption in N2 control and *ifb-1(ju71)* worms. (D) Oxygen consumption over the assayed period. (E) Representative images of nematodes expressing *Posm-5::mito::gfp* stained with (mitochondria membrane potential sensitive) TMRE dye in different genetic backgrounds. (F) ICQ quantification of mito::GFP and TMRE staining in different genetic backgrounds (N = 15 worms). Scale bar: 10 μm. Error bars: ± max. and min. range. N = 50 worms per well with 10 wells/study group (A-D). Student’s t-test. *p<0.05, **p<0.01.

We then examined colocalization between mitochondria and overexpressed IFB-1 in *C. elegans* amphids. Here, we coinjected a *Posm-5::mito::gfp* construct with a *Posm-5::ifb-1a::mrfp* construct into N2 worms. Figure 6A-D shows the head region of the worm, similar as shown in Figure 1 (H+J). Cell bodies of amphid neurons (located in the nerve ring) are clearly visible to the right from which the dendritic extension of amphid neurons emanate to the tip (“nose”) of the worm located to the left (Fig. 6A). At this (rather low) magnification distinct IFB-1 particles (or cluster) can be seen in somas (Fig. 6B), but barely detected in the dendritic bundles. Thus, we also provide higher magnification images (Fig. 6E), revealing single mitochondria (arrows), IFB-1 aggregates (big arrowhead), or both particles colocalized (small arrow heads). Figure 6C is a merged image of Figure 6A+B showing various colocalization events in somas and Figure 6D is a false color representation of PDM values (more positive values reflect more accurate colocalization, see also Methods). Lastly, we expressed IFB-1A::GFP in vimentin knockout cells (SW13) and used differential centrifugation to prepare a crude mitochondrial fraction. Critically, we were able to detect IFB-1 protein in these fractions. As a control, we expressed GFP protein alone which was not detected in the mitochondria fraction (Fig. 6G).

**Figure 6:**
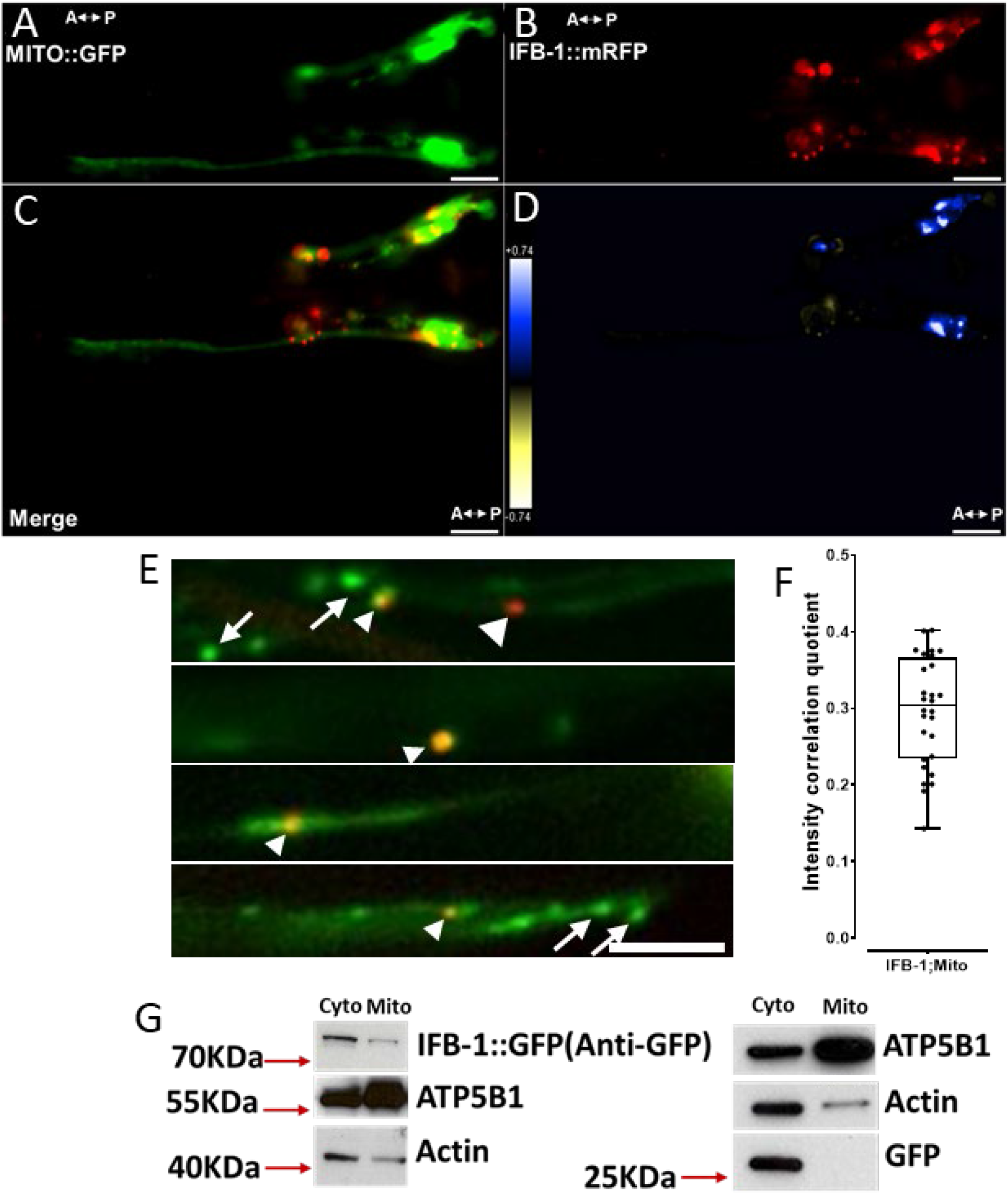
Mitochondria/IFB-1 colocalization and physical interactions between mitochondria and IFB-1 protein. (A-D) Images of wild type worms co-expressing mito::GFP and ifb-1::mRFP under the Posm-5 promoter. (E) Magnified images with short arrows pointing to specific colocalization events, arrow heads pointing to exemplified IFB-1 particles and long arrows pointing to exemplified single mitochondria. (F) ICQ quantification of IFB-1/mitochondria colocalization from (C). (G) Western blot of crude mitochondrial fraction (Mito) and cytosolic fraction (Cyto) isolated from SW13 (vim-) cells expressing either IFB-1::GFP (left panel) or only GFP (right panel). ATP5B1 antibody used to detect solubilized mitochondria. Scale bars: 20 μm in (A-D) and 5 μm in (E). Error bars: ± max. and min. range. N = 28 worms. A – anterior direction and P – posterior direction.

## DISCUSSION

Abnormal mitochondria distributions and neuronal IF accumulations is a hallmark of various neurological diseases ^1,3,22^. Especially neurons are cells with high energy demand based on their dedicated functions such as synaptic activities. Consequently, mitochondria are of critical importance and not only pathological dysfunction of mitochondria (based on genetic defects or environmental clues such as ROS), but also the role of mitochondrial dynamics (fusion and fission events, transport efficiencies) in neurodegeneration are widely discussed ^22,39–41^. Thus, proper and balanced axonal transport of mitochondria is critical for neuronal function and viability. On the other hand, only little is known how mitochondria transport and distribution is regulated. Here, we report that IFB-1 is a crucial factor for regulating the balance of mitochondrial transport, and thus, distribution (Figs. 2+3).

As mentioned above, the basic molecular mechanisms how mitochondria are transported are known; however, mechanisms that lead to the immobilization of mitochondria (specifically at sites of high energy demand) are less investigated. Kinesin-1 (the major transporter of mitochondria in neurons) can be regulated by kinase activities (e.g., PI3-kinase, CDK5 or GSK3) triggering the motor’s stops and starts, or specific adaptors (receptors) such as the Miro/Milton (Trak) complex to regulate the binding of mitochondria to the motor (depending on local calcium levels) ^22,42–44^. Vice versa, static anchors such as the microtubule binding protein SNPH actively stop mitochondria at ATP-deficient sites ^27^. Also NFs, desmin and vimentin all bind to mitochondria regulating their dynamics and distributions ^9,26,32,33,45,46^. But also IF-associated proteins such as plectin 1b are able to connect the IF-network to mitochondria ^35^.

*C. elegans* chemotaxis behavior is largely regulated by the amphid chemosensory organs which are paired bundles of sensory neurons laterally running along the head of the worm (Figs. 1A and H-J, and 6A-D). Each amphid bundle contains 11 ciliated chemosensory neurons (and one thermosensory neuron) all sensing a characteristic set of attractants, repellents and pheromones. Somas and (relatively short) axons are both located in the nerve ring from where long dendrites emanate to the tip of the nose where their ciliated endings are exposed to the environment ^38,47^. In most mammalian neurons, microtubule polarity is mixed in dendrites, however, dendrites of *C. elegans* amphid neurons are arranged uniformly, similar to the microtubule arrangement in *D. melanogaster* sensory neurons ^48^. Thus, anterograde and retrograde directions of moving mitochondria can be accurately distinguished. The velocities of mitochondria in these neurites (Figs. 2 and 4) are comparable to velocities recorded in mammalian systems ranging from 0.32 to 0.91 μm/s. Similarly, the mobile pool (processive + vibrating mitochondria) was determined to be 50% in *C. elegans* amphid neuron bundles (Fig. 4E), a number close to that reported in mammalian systems (>65%) ^49^. Note that we provide rescue experiments for most motility data in *ju71* mutants (Fig. 2) underscoring significance and accuracy of our findings. The general effect of IFB-1 suppression on mitochondrial dynamics was reduced speeds in both anterograde and retrograde directions immediately resulting in less reversals (Fig. 2) and increased number of stationary mitochondria (Figs. 2A-C and 4F). These effects result in unbalanced mitochondria distributions with reduced densities but increased cluster sizes (Fig. 3).

We thus hypothesize that transient interactions of mitochondria with the IFB-1 network is critical for balanced organelle movements and distributions. To understand whether depletion of ifb-1 may affect the fusion/fission machinery (or vice versa), we knocked down genes fis-1 and eat-3 (OPA-1 homolog) in N2 and *ju71* mutants. Effects were mostly as expected in N2 (except that densities were unaffected after eat-3 knockdown) and it is interesting to observe that fis-1 knockdown had additive effects on *ju71* while eat-3 knockdown did not further change the ifb-1 phenotype. These results may point to additive effects in parallel pathways between fis-1 and ifb-1 genes and independent eat-3 and ifb-1 pathways. In our model (Fig. 7), the IF network provides additional support and transient anchor points for large mitochondria during their movements on microtubules. We suspect that without these additional anchor points, mitochondria movement would be less smooth resulting in frequently microtube detachments, off-axis movements and microtubule switching. Such unbalanced transport not only results in mitochondria accumulations (Fig. 3) but also in impaired metabolism directly affecting oxygen consumption of nematodes (Fig. 5); as well as leading to developmental defects (such as shorter neurites and axons, Fig. 1K+M) with the consequence of impeded chemosensation (Fig. 1C+D). It is critical to note that in our model mitochondria interact with the IF network only transiently: during their movement on the microtubule network mitochondria dynamically attach and detach from specific IF anchor points. This assumption is in accordance with colocalization experiments showing that mitochondria are not always in close proximity to IFB-1 cluster (Fig. 6E), whereas IFB-1 protein is generally able to physically interact with mitochondria (Fig. 6G). Therefore, while IFs might serve as enduring docking sites at localized areas of neurons (with high energy demand), they may also act as transient (short-termed) attachment points for these large organelles transported by comparably minuscule molecular motors (likely not being able to fully stabilize mitochondria on microtubules). Indeed, it has been suggested that the transport of large vesicular structures along microtubule tracks (in the thin and overcrowded axon, ^50^) would find additional support at opposing actin tracks just below the plasma membrane in axons (actin cortex) ^31,42,49^. Notably, Leterrier *et al*. 1994 published unique electron micrographs of mitochondria docking to NFS and that Wagner *et al*. 2003 used atomic force microscopy (as well as TEM) to reveal direct interactions between mitochondria and NFs ^26,51^. We also propose that surface charges may be responsible for such transient mitochondria/IF interactions (Fig. 7). Here, the high membrane potential of mitochondria may facilitate the interaction with the negatively charged IFB-1 protein (net charge is −11.6 in cytoplasm with an isoelectric point of 5.43, isoelectric.org). In more detail, the head domain of IFB-1 has an isoelectric point of 6.71 and a charge of −0.1 in the cytoplasm, the rod domain inherits an isoelectric point of 5.03 and a charge of −13 in the cytoplasm while the tail domain of IFB-1 has an isoelectric point of 8.35 with a charge of +2.4 (isoelectric.org).

**Figure 7:**
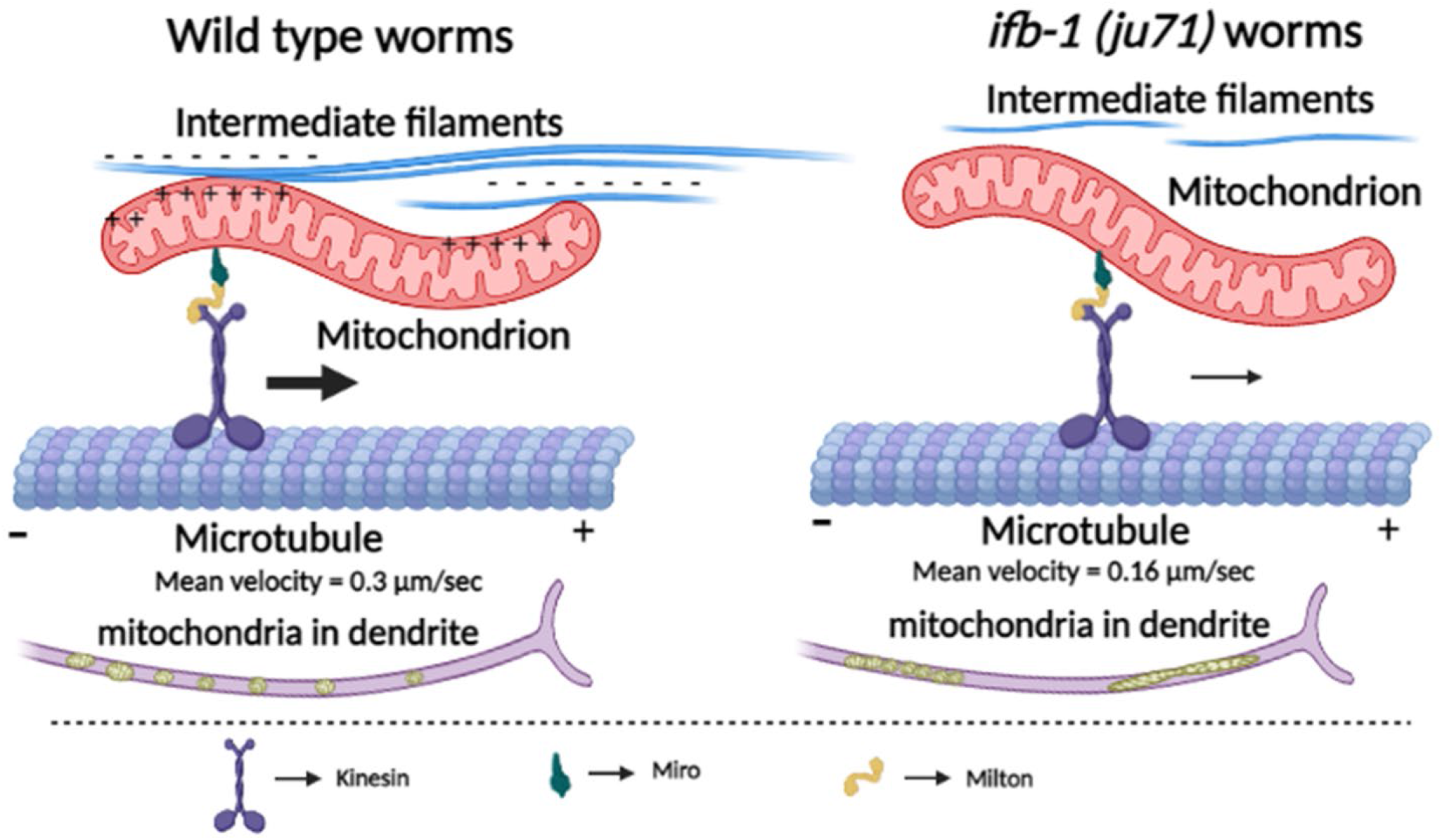
Model summarizing this study. To the left: Additional (transient) mechanical support based on charge interactions provided by IFs for the long-range transport of (large) mitochondria by (relatively small) kinesins. To the right: In IFB-1 mutants, the interaction between mitochondria and IFs is decreased resulting in imbalanced transport and thus accumulations in dendrites of amphid sensory neurons.

## MATERIAL AND METHODS

### Worm maintenance, strains and plasmids

Worms were maintained at 20°C on NGM agar plates seeded with OP50 *E. coli* according to standard methods ^52^. Strains CZ4380 *ifb-1(ju71)* and RB1253 *ifb-1(ok1317)* were received from the CGC (Caenorhabditis Genetics Center, Minnesota, USA). The *ju71* allele carries a 617 bp deletion that removes the promoter region of *ifb-1a* (Fig. 1N) abolishing the expression of IFB-1A (but not IFB-1B) protein based on Western blots ^18,37^. The *ok1317* has not been characterized yet and based on our analysis, it carries a 1361 bp deletion located upstream of the *ju71* deletion, yet residing in the *ifb-1a* promoter region. Both alleles (*ju71* and *ok1317*) likely remove parts of promoter in the ifb-1a gene likely affecting its expression level (Fig. 1G+N). In order to visualize mitochondria in *C. elegans* amphid sensory neuron bundles, we employ the *Posm-5* promoter known to selectively express genes in amphid, labial and phasmid neurons ^16,17^. Here, we used an existing construct *Posm-5::xbx-1::yfp* ^53^ and replaced the *xbx-1::yfp* insert with an *mito::gfp* insert (encoding for a C-terminal GFP fused to an N-terminal mitochondria-targeting sequence) amplified from the Andrew Fire vector pPD96.32 (Plasmid #1504, Addgene, Cambridge, MA). The resulting plasmid *Posm-5::mito::gfp* was coinjected with the roller marker *pRF4(rol-6(su1006))* into N2 worms (100 ng/μl) using standard microinjection techniques ^54,55^. *N2;nthEx4[Posm-5::mito::gfp;rol-6(su1006)]* worms were then crossed with the aforementioned strains CZ4380 and RB1253, respectively, and genotyped for homozygosity using following primers: For the *ju71* mutation 5’-CCGATTTATTGCAAGATGGTGTAGA-3’ (forward), 5’-TCAATGGTTTTGCCTTCCCG-3’ (reverse) and for the *ok1317* mutation 5’-AATCAGCGCCACAGGC-3’ (forward) and 5’-GCAACCAATTTACGGTTCTG-3’ (reverse). The rescue strain was obtained by microinjecting the plasmids *Posm-5::ifb-1::mrfp*, *Posm-5::mito::gfp* along with the roller marker pRF4*(rol-6(su1006))* into *ifb-1(ju71)* worms. For colocalization experiments, we constructed a *Posm-5::ifb-1a::mrfp* vector by subcloning the *ifb-1* insert from and existing vector pCZ454 *Pifb-1::ifb-1a::gfp* (a kind gift from Dr. Andrew Chisholm) and the *mrfp* from an existing vector *Punc-104::snb-1::mrfp* ^56^. *Posm-5::ifb-1a::mrfp* was then coinjected with *Posm-5::mito::gfp* (at both 100 ng/μl) and pRF4*(rol-6(su1006))* (50 ng/μl) into N2 worms to obtain the strain *N2;nthEx5[Posm-5::ifb-1a::mrfp;Posm-5::mito::gfp;rol-6(su1006)]*. Complementary DNA of *ifb-1* was cloned into expression vector pEGFP-N3 (a kind gift from Prof. Yin-Chang Liu, NTHU, Taiwan) using AAAAGCTAGCATGGAGGCTGAATCAGC (forward) and AAAAGGTACCTTGTCCGGATTGGATGG (reverse) between NheI and KpnI restriction sites for transfection into SW13 (vim-) cells.

### Dye-filling and chemotaxis quadrant assay

To investigate dye uptake of sensory neurons, we incubated worms in M9 buffer containing a final concentration of 0.1 mg/ml DiI (1,1’-dioctadecyl-3,3,3’3’-tetramethylindo-carbocyanine perchlorate, Sigma Aldrich) in 0.5% Triton X-100 ^57^. After 2 hours incubation on a rotator, worms were subsequently mounted on 2% agarose pads and anesthetized with 5 mM tetramisole (Sigma-Aldrich) for observation under a fluorescence microscope.

Standard quadrant assay to test chemotaxis of worms was performed as described by Margie et al. ^58^. NGM plates (5 cm, without OP50) were prepared according to the protocol and 100 to 200 synchronized worms at the young adult stage were transferred to the center of each plate. Plates were inverted and incubated at room temperature for 1 to 2 hours. After incubation, worms at different quadrants were counted immediately. 2 μl of chemoattractant (0.5% diacetyl mixed with equal volume of 0.5 M sodium azide) and control solution (0.5 M sodium azide mixed with equal volume of ddH_2_O, the solvent for diacetyl) were placed at designated positions (Fig. 1C) just before the addition of worms. Chemotaxis index is calculated by: CI = (number of worms in test quadrants)-(number of worms in control quadrants)/total number of worms.

### Mitochondria motility, particle size and colocalization analysis

To record the movement of mitochondria in amphid sensory neuron bundles, we immobilized living nematodes using 5 mM tetramisole (Sigma-Aldrich) before placing them on 2% agarose-coated objective slides. An Olympus IX81 microscope with a DSU Nipkow spinning disk unit connected to an Andor iXon DV897 EMCCD camera was employed for high-resolution and high-speed time-lapse imaging (3-4 frames per second). To create kymographs from recorded videos, we used the image processing and analysis software NIH ImageJ (NIH, http://rsb.info.nih.gov/ij/). Here, we draw a line over a neuron of interest (within the video) and after applying the “reslice stack” function, a stack of single sequences from the movie at the defined area is generated. Static particles then appear as vertical lines while moving particles leave traces on horizontal directions (e.g., Fig. 2E), and from the slopes of these traces velocities of these particles can be calculated. Curved neurons were straightened before their conversion into kymographs using the “straighten curved objects tool”. A moving event is defined as a single motility occurrence typically right after a pause or a reversal and, vice versa, it ends when the particle is again pausing or reversing. Mitochondrial moving distances larger than 1.6 μm were classified as “processive”, otherwise as “vibrating” (see also ^45^). For mitochondria particle analysis such as area, Feret’s diameter and density (Fig. 3), we used the “particle analysis” plugin from ImageJ. For neurite length (Fig. 1K+M), we used the “line measurement tool”. For colocalization analysis, the “intensity correlation analysis” plugin from ImageJ was employed according to a method developed by Li et al. ^59^ (Fig. 5F+6F). The intensity correlation quotient (ICQ) is based on the PDM (<u>p</u>roduct of the <u>d</u>ifferences from the <u>m</u>ean) that is equal to: (red intensity – mean red intensity) × (green intensity – mean green intensity). ICQ values range from −0.5 to 0.5 with values close to 0.5 standing for interdependent expression of two fluorophores, close to −0.5 for segregated expression and values near 0 for random expression of two fluorophores.

All investigated worms in this study are at young adult stages.

### RT-qPCR

For RT-qPCR (reverse transcription quantitative PCR), we designed the following primers that cover the 1st and 3rd exon (229 bp) of the *ifb-1a* gene: 5’-AATCAGCGCCACAGGC-3’ (forward) and 5’-CAACCAATTTACGGTTCTG (reverse). As an internal control, we use *cdc-42* gene ^60^ with the following primer set: 5’-CTGCTGGACAGGAAGATTACG-3’(forward) and 5’-CTCGGACATTCTCGAATGAAG-3’(reverse). To quantify PCR products, we use the ABI Power SYBR Green PCR Master Mix (ABI, USA) in conjunction with an ABI 7500 real-time PCR machine.

### Whole worm oxygen consumption analysis

Oxygen consumption analysis was performed according to a method described by Yamamoto et al. ^61^ employing the XF24 Extracellular Flux Analyzer (Seahorse Bioscience Inc., USA). Worms were synchronized to obtain young adults for analysis and washed several times before placing them on NGM agar plates without OP50. 50 worms were then picked to each vial of a standard 24-well Seahorse plates (#100777-004) containing 200 μl of M9 buffer. Four of these 24-wells were used for background correction (only buffer) and all remaining wells were divided equally for two different experimental groups (N2 and *ifb-1(ok1317)* or N2 and *ifb-1(ju71)*). Oxygen consumption by the worms from the medium was measured 15 times over a period of 120 min in real-time. Data from different experimental groups were analyzed using Student’s t-test.

### RNAi feeding

NGM plates for RNAi feeding experiments were prepared with ampicillin (100 μg/ml) and 1 mM IPTG. Plates were spread with HT115 *E. coli* strains carrying dsRNA against the fis-1 and eat-3 genes and incubated overnight at 37°C. 10 worms expressing mito::GFP were added to the fis-1 and eat-3 RNAi plates and transferred to new plates every day. Every 24 hrs, worms were washed to fresh RNAi plates. After 48 hrs of feeding on RNAi plates worms were immobilized using 5 mM tetramisole (Sigma-Aldrich) before placing them on 2% agarose-coated objective slides and subjected to imaging for mitochondrial cluster analysis.

### Life span assay

The life span assay protocol described by ^62^ was slightly modified, where we used age synchronized worms by timed egg laying on a plate for 2-3 hrs. Worms were then transferred to newly prepared NGM plates with heat-deactivated ^63^ (65°C for 15 min) OP50 *E. coli* and transferred to new plates every day. Dead worms were identified by their motionlessness when prodded with a platinum wire. These worms were then scored excluding those that disappeared or died by otherwise causes.

### Worm neuronal culture

Isolation and culturing primary neurons form *C. elegans* embryos was performed based on protocols by Christensen et al. ^64^ and Strange et al. ^65^. Isolated neurons are maintained in Leibowitz’s (L15) medium overlaid with 10% fetal bovine serum (FBS), 100U/ml of penicillin, 100 μg/ml of neomycin and 100 μg/ml of streptomycin. These cultures can be stored for up to 10 days at 20°C without the need of adjusting the CO_2_ atmosphere.

### Mammalian cell culture

Mammalian cell line SW13 (vim-) was a kind gift from Prof. Ming-Der Perng (National Tsing Hua University, Taiwan). SW13 (vim-) cells were cultured in DMEM medium with 10% fetal bovine serum at 37°C with 5% CO_2_ atmosphere. At 70-80% confluency, cells were transfected with either plasmid construct pEGFP-N3-IFB-1 or pEGFP-N3 (without insert as a control) using K2^®^ transfection system from Biontex (Germany).

### Mitochondrial pull down assay and Western blotting

We used SW13 vim-cells to overexpress IFB-1 and to obtain a crude mitochondria fraction. Here, we washed SW13 cells with PBS thrice to remove culture media, trypsinized them, resuspended in isolation buffer and homogenized on ice using a glass homogenizer (20 to 25 strokes). Homogenized solution was then centrifuged at 1000 x g at 4°C for 10 min. The supernatant was collected and centrifuged at 14,000 x g for 15 min at 4°C gaining a cytosolic fraction with the pellet containing crude mitochondria ^66^. The pellet was then washed twice with isolation buffer and both cytosolic and mitochondrial fractions were quantified for its protein concentration using Thermo Scientific Pierce^™^ BCA protein assay kit (#23225), whereas 30 μg of protein was used for Western blotting analysis. 8% SDS-PAGE was used for protein separation by electrophoresis and transferred to a PVDF membrane (Immobilon^®^-P, Millipore Corporation, IPVH00010). 10% BSA was used to block the membrane followed by protein detection using primary antibody of 1:1000 dilution such as anti-β actin (C4, SC-47778, mouse monoclonal antibody, Santa Cruz Biotechnology, USA), anti-GFP (B-2, SC-9996, mouse monoclonal, Santa Cruz Biotechnology, USA) and ATP5B1 (Gene Tex Inc.), respectively; as well as secondary antibody of 1:5000 dilution in conjunction with Luminata^™^ Classico Western HRP substrate (Millipore Corporation, WBLUC0500).

### TMRE staining

TMRE staining of mitochondria in *C. elegans* was performed as described in ^67^ with little modification. Young adult worms were washed with M9 buffer to remove the bacteria. TMRE dye (a kind gift from Prof. Chuang-Rung Chang, NTHU, Taiwan) with a final concentration of 300 nM in M9 was added and the solution rotated for 8 hours at 20°C in the dark. The worms were then washed four times with M9 buffer and were placed on OP50 seeded NGM plates for 1-2 hours in the dark at 20°C. Worms were then mounted onto 2% agarose pads for imaging.

## Supporting information

mut video

## ACKNOWLEDGEMENTS

We thank the *C. elegans* Core Facility (CECF) Taiwan funded by National Science Council (NSC) for providing us Andrew Fire Vectors, microinjection setups and worm observation systems. We also thank lab members Yu-Fei Peng, Chih-Wei Chen and Víctor Aplícano for their assistance. We further thank colleagues Dr. Yin-Chang Liu, Dr. Horng-Dar Wang, Dr. Chuang-Rong Chang and Dr. Mou-Chieh Kao for assistance. We thank Aravind Chandrasekaran for assisting with movie conversions. This work was funded by NSC grants NSC 100-2311-B-007-004 and 101-2311-B-007-002-to OIW.

## AUTHORS’ CONTRIBUTIONS

SNB - Designed and performed experiments, analyzed data and wrote the manuscript; MMS - Designed and performed experiments, analyzed data and wrote the manuscript; YC – Designed and performed experiments, analyzed data and wrote the manuscript; PB – Performed experiments and analyzed data; GHW - Performed experiment and analyzed data; OIW – Designed experiments, obtained grants and wrote the manuscript.

---

**Supplemental Video 1**: Video sequence of recorded mitochondria movements in amphid sensory neurons of (A) wt and (B) *ju71*. Time-lapse factor: 4 x (equals to 28 frames per second).

